# Optimizing non-invasive sampling of an infectious bat virus

**DOI:** 10.1101/401968

**Authors:** John R. Giles, Alison J. Peel, Konstans Wells, Raina K. Plowright, Hamish McCallum, Olivier Restif

## Abstract

Notable outbreaks of infectious viruses resulting from spillover events from bats have brought much attention to the ecological origins of bat-borne zoonoses, resulting in an increase in ecological and epidemiological studies on bat populations in Africa, Asia, and Australia. The aim of many of these studies is to identify new viral agents with field sampling methods that collect pooled urine samples from large plastic sheets placed under a bat roost. The efficiency of under-roost sampling also makes it an attractive method for gathering roost-level prevalence data. However, the method allows multiple individuals to contribute to a pooled sample, potentially introducing positive bias. To assess the ability of under-roost sampling to accurately estimate viral prevalence, we constructed a probabilistic model to explore the relationship between four sampling designs (quadrant, uniform, stratified, and random) and estimation bias. We modeled bat density and movement with a Poisson cluster process and spatial kernels, and simulated the four under-roost sheet sampling designs by manipulating a spatial grid of hexagonal tiles. We performed global sensitivity analyses to identify major sources of estimation bias and provide recommendations for field studies that wish to estimate roost-level prevalence. We found that the quadrant-based design had a positive bias 5–7 times higher than other designs due to spatial auto-correlation among sampling sheets and clustering of bats in the roost. The sampling technique is therefore highly sensitive to viral presence; but lacks specificity, providing poor information regarding dynamics in viral prevalence. Given population sizes of 5000–14000, our simulation results indicate that using a stratified random design to collect 30–40 urine samples from 80–100 sheets, each with an area of 0.75–1m^2^, would provide sufficient estimation of true prevalence with minimum sampling bias and false negatives. However, acknowledging the general problem of data aggregation, we emphasize that robust inference of true prevalence from field data require information of underpinning roost sizes. Our findings refine our understanding of the under-roost sampling technique with the aim of increasing its specificity, and suggest that the method be further developed as an efficient non-invasive sampling technique that provides roost-level estimates of viral prevalence within a bat population.

## Introduction

Recent emergence of bat-borne viruses has motivated an increase in ecological and epidemiological studies on bat populations in Africa, Asia, and Australia (Calisher et al. 2006, Halpin et al. 2007, Wang and Cowled 2015). Little was known about these pathogens at the outset of investigation, so research focused first on discovering the reservoir host(s), as demonstrated by Hendra virus in Australia (Halpin et al. 2000), Nipah virus in Malaysia (Chua et al. 2002), SARS in China (Li et al. 2005), Marburg in Africa (Towner et al. 2009), and Ebola viruses in Africa (Breman et al. 1999) and the Philippines (Jayme et al. 2015). Further, considerable effort is now invested into identifying additional unknown viral pathogens in bats that have epidemic potential; an important undertaking that minimizes spillover risk via vaccine development, predicting epidemic potential, and developing assays to detect the virus in humans and wildlife (Anthony et al. 2013, Drexler et al. 2012, Quan et al. 2013, Smith and Wang 2013). Discovering a virus and identifying its reservoir host(s) is also the first step in describing viral *dynamics* (patterns of viral presence in bat populations over space and time), which provide insights into the broader ecological context surrounding spillover and precursors to the emergence of bat-borne viral diseases in humans (Hayman et al. 2013, Plowright et al. 2015, Wood et al. 2012).

A common approach, in bat-borne disease research, involves the capture of many individual bats repeatedly over time, where bats are sampled (e.g. serum, urine, saliva) and tested for viral presence using serology or PCR techniques. Best case scenario, longitudinal samples are obtained for multiple individuals, enabling both the discovery of new viruses and description of dynamics in individual-level viral prevalence. Individual-level longitudinal data are more common for high-fidelity cave-roosting bats which can be recaptured frequently at the same roosting site (Streicker et al. 2012, Towner et al. 2009). However, these type of longitudinal data are much more difficult to gather from the tree roosting megachiroptera, such as Pteropus and Eidolon genera (Hayman et al. 2012), which are highly-mobile nomadic foragers, making them poor candidates for ecological studies that rely on recapture of individuals. Therefore, recent research has supplemented the capture of individual bats with a non-invasive sampling technique that uses plastic sheets to collect urine and feces under bat roosts (Baker et al. 2012, Baker et al. 2013, Chua 2003, Chua et al. 2002, 2001, Edson et al. 2015a, Field et al. 2011, 2015, Marsh et al. 2012, Pritchard et al. 2006, Smith et al. 2011, Wacharapluesadee et al. 2010).

For several viruses of public health interest, urinary excretion is a primary route of transmission (e.g. Nipah virus in Asia and Australia (Middleton et al. 2007, Wacharapluesadee et al. 2005), Hendra virus in Australia (Edson et al. 2015b), and both Henipaviruses (Iehlé et al. 2007) and Marburg virus in Africa (Amman et al. 2012)). The under-roost sampling technique takes advantage of this particular mode of transmission to achieve longitudinal sampling of a bat population at the *roost-scale* that is both cost-effective and reduces exposure to infectious viruses compared to catching individual bats. Under-roost sheet sampling was initially implemented in 1998 to isolate Nipah and Tioman virus from *Pteropus hypomelanus* and *P. vampyrus* in Malaysia (Chua 2003, Chua et al. 2002, 2001). Under-roost sampling designs typically use large sheets placed under roost trees, and urine droplets are pooled into an aggregate sample from the area (or sub-area) of each sheet. Most studies provide minimal description of the sheet sampling design, however Edson et al. (2015a), Field et al. (2015), and Wacharapluesadee et al. (2010) describe their methods in greater detail (i.e. sheet dimensions, number of sheets, pooling of urine samples). In general, the under-roost sampling technique was initially designed to isolate viral agents, not necessarily study viral dynamics, however a few recent studies have also employed the technique to collect longitudinal data and describe patterns in viral prevalence for Nipah virus in Malaysia (Wacharapluesadee et al. 2010) and Hendra virus in Australia (Field et al. 2015, Páez et al. 2017). However, the extent to which the data are vulnerable to sampling bias has not been explored.

The most salient complication is that under-roost sampling estimates individuallevel prevalence with sheet-level prevalence. In this scenario, binomial samples are comprised of urine droplets from an ‘area’, which are pooled to constitute sufficient volume for an array of molecular assays (i.e. PCR and/or whole genome sequencing). Although this is a necessary compromise, the clustered nature of bat density within a roost acts as a confounder that allows an unknown number of individuals to contribute to a sample. In this manner, under-roost sampling may introduce systematic sampling bias in the form of increased sensitivity of viral detection assays.

The increased sensitivity of pooled samples is well-known. Sample pooling was first used during world war II to avoid the ‘expensive and tedious’ process of monitoring syphilis in US soldiers (Dorfman 1943), and since, it has been used as a cost-effective method to screen for HIV infection in developing countries (Behets et al. 1990). ‘Herdlevel’ testing is also common in surveillance of livestock diseases where a pooled sample is used to determine presence or absence of a disease within the herd (Christensen and Gardner 2000); if the herd is found positive, individual-level samples are then used to identify infected individuals or calculate prevalence more accurately (Litvak et al. 1994). In this regard, pooling urine samples as part of the under-roost sampling method is well-suited for surveillance of bat viruses because the higher sensitivity of pooled sample testing is advantageous when individual-level prevalence is very low (Muñoz-Zanzi et al. 2006). Conversely, the high sensitivity of pooled samples is problematic when used to estimate individual-level prevalence (Cowling et al. 1999)—a classic statistical problem resulting from data aggregation, often referred to as the ‘ecological fallacy’ (Robinson 2009).

Our aim, therefore, is to contribute the first modeling study to theoretically explore the application of under-roost sheet sampling in a generic tree roosting bat population and quantify the potential sampling bias introduced by different sampling techniques. We focus on tree roosting pteropid bats because they are reservoir hosts of several viruses considered to be a public health risk, and based on their highly mobile population structure, under-roost sampling techniques are especially useful. Specifically, we explore four questions in detail: 1) Given different under-roost sheet sampling designs, how accurately is individual-level viral prevalence estimated? 2) What is the estimation bias across all values of individual-level prevalence? 3) What are the major drivers of estimation bias? And 4) If you reduce the size of the sheets on which samples are pooled, and increase their number, can you reduce sampling bias and provide an acceptable estimate of individual-level prevalence? To address these questions, we designed four simulation scenarios comprised of a probabilistic model of bat density within a generic roost of tree roosting pteropid bats and four under-roost sheet sampling designs (quadrant, uniform, stratified, and random). We then explore the parameter space of these scenarios and perform global sensitivity analysis to determine the primary drivers of estimation bias. Our results provide some useful recommendations on how to apply under-roost sampling for the surveillance of infectious bat viruses.

## Methods

### Modeling bat density in a roost

Pteropid bat roosts can be spread out and encompass many trees, with individuals moving frequently within the roost, so we modeled bat density within a generic bat roost with a Poisson cluster process of roosting positions and a spatial Gompertz probability density function that reflects movement within a roosting site. Specifically, bat density within roost area *A* (a disc with radius *r*) is constructed in four stages that include: 1) placement of roosting trees within the roost area, 2) clustering of individuals around them, 3) individual-level movement within a tree, and 4) a separate model of roost-wide movement. We used a Thomas cluster process to simulate the spatial clustering of bat positions around trees, using the `rThomas` function from the `spatstat` package in the R programming language (Baddeley et al. 2015, R Core Team 2016). Tree locations (parent points) were randomly distributed within *A* subject to a homogeneous intensity *κ*, given by *n*_*t*_*/A*, where *n*_*t*_ is the number of occupied trees in the roost. The mean number of bats in each roost tree *µ* is simulated by the cluster point process which is Poisson distributed with mean *µ*. Individual bat positions are determined according to an isotropic Gaussian kernel centered on each tree with radius *r*_*t*_. Note that even when parameters *κ, r*_*t*_, and *µ* are fixed, the number of bats in the roost *N*_*b*_ will still vary upon each simulation because the Poisson point process is stochastic.

Bat movement was modeled at the individual-level and roost-level. To model individual-level movement, we calculated a kernel density estimate for the simulated point process that sums Gaussian kernels with a radius of 0.5m centered on each bat position. We modeled roost-wide movement with a spatial Gompertz probability density using the `dgompertz` function from the `flexsurv` package (Jackson 2014). The distribution of the Gompertz is controlled by shape and rate parameters that determine the function’s curvature and rate of decay respectively. We chose ranges for the these parameters that make the least assumptions about movement, where values are high for a large area at the roost’s center, but decay quickly toward the edges. To make the final kernel density estimate for bat density, we combined models of individual- and roost-level movement and ensured that the function integrated to 1 (Figure 1).

**Figure 1:**
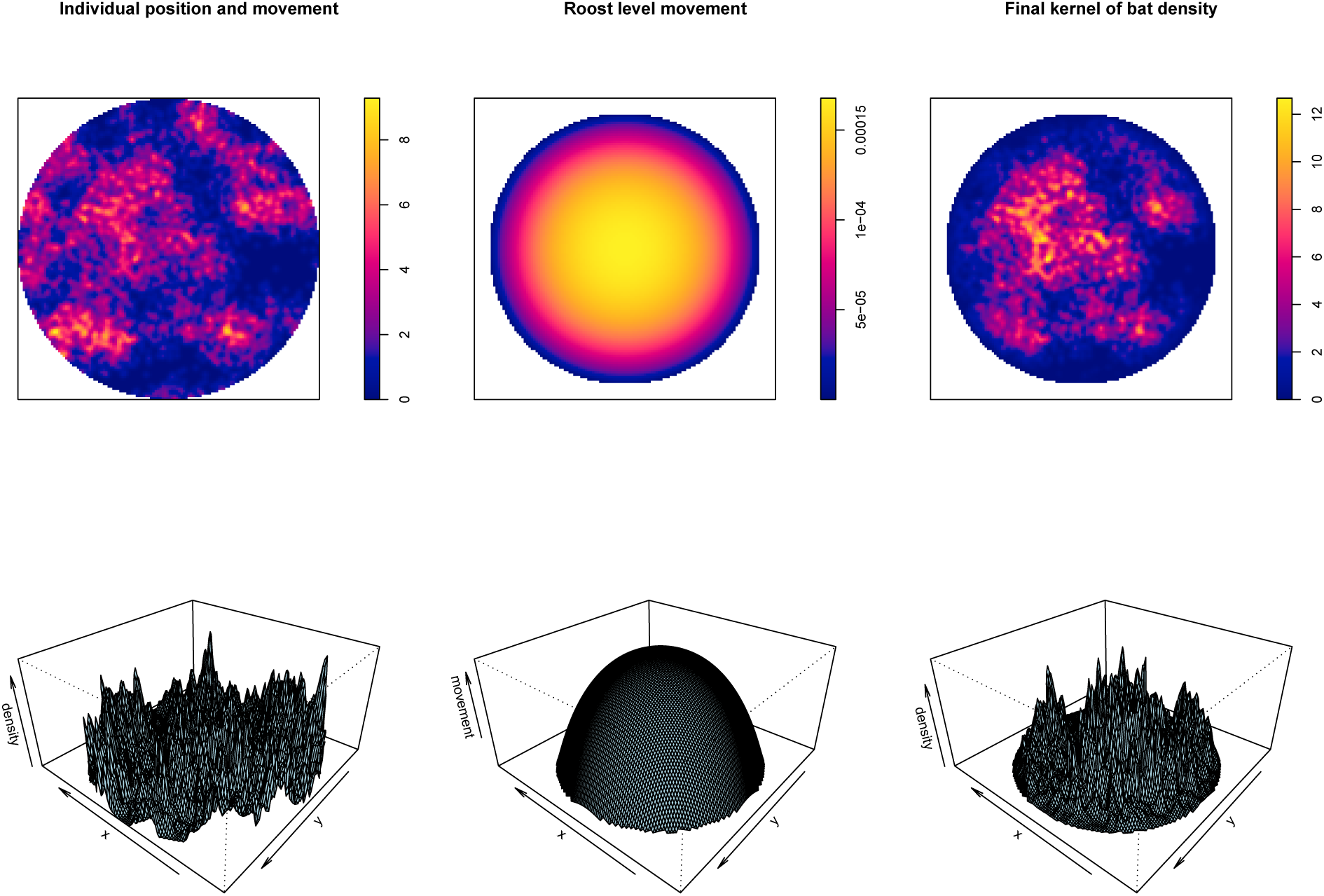
Illustration of one simulation of a kernel density estimation of bat density within a roost. The top row shows pixel-images and the bottom row shows perspective plots of: the density of roosting positions and individual-level movement around them (left), an isometric Gompertz probability density function centered on the roost to model roost-level movement (middle), and the final estimated intensity function used to model bat density (right).

### Modeling under-roost sheet sampling

We explored the effect of four different under-roost sheet sampling designs: quadrant, uniform, stratified, and random. An efficient way to simulate each sampling design within two-dimensional circular space uses hexagonal tiles, where the size and combination of tiles selected can replicate different sheet-based sampling designs. We calculated the number of bats roosting and moving above a sampling sheet by using the area of each hexagonal polygon to define the space of integration *S*.

We determined the dimensions for the quadrant-based design using common protocols for under-roost sheet sampling of Australian fruit bats found in Edson et al. (2015a) and Field et al. (2015). Here, 10 large 3.6 × 2.6m sheets were placed under the roost and divided into 1.8 × 1.3m quadrants, where urine samples were pooled within each quadrant (allowing up to 4 samples per large sheet). Considering each quadrant to be its own ‘sheet’, we replicated this sampling design by making a hexagonal grid with each tile area equivalent to a 1.8 × 1.3m rectangular sheet. Groupings of 4 hexagonal tiles then suffice as a large sheet with 4 quadrants. In each simulation, we generated 10 sheet positions within *A* using a simple sequential inhibition point process with the `rSSI` function of the `spatstat` package (Baddeley et al. 2015). To ensure that all sheets retained the same quadrant orientation and that no two sheets were directly adjacent, we generated sheet positions within a disc of *A* − 3m and set the inhibitory radius to 3*s*, where *s* is the hexagonal cell size. The four cell-centers nearest each of the 10 simulated point locations comprised the 40 (10 × 4 quadrants) hexagonal tiles for the quadrant-based design (S1).

To test our hypothesis that a larger number of smaller sheets will estimate roost-level prevalence more accurately, we generated hexagonal grids with cell size *s* that select *h* number of tiles in a uniform, stratified, or random pattern. Both uniform and random designs are straightforward, but the stratified sampling design was generated using a sequential inhibition point process, where random points are laid down sequentially, retaining only those that are placed further than a specified inhibitory radius *r*_*s*_. This is similar to a person attempting to lay down sheets randomly with one rule in mind—“Do not place sheets within *r*_*s*_ distance of each other”. We simulated sheet sampling designs with the `sheetsamp` function in the R code provided in Supplementary Information. Figure 2 displays an example of a simulation which has generated the previously implemented large-sheet quadrant design and three additional ‘small-sheet’ designs that use a larger number of smaller (1 × 1m) more dispersed sheets.

**Figure 2:**
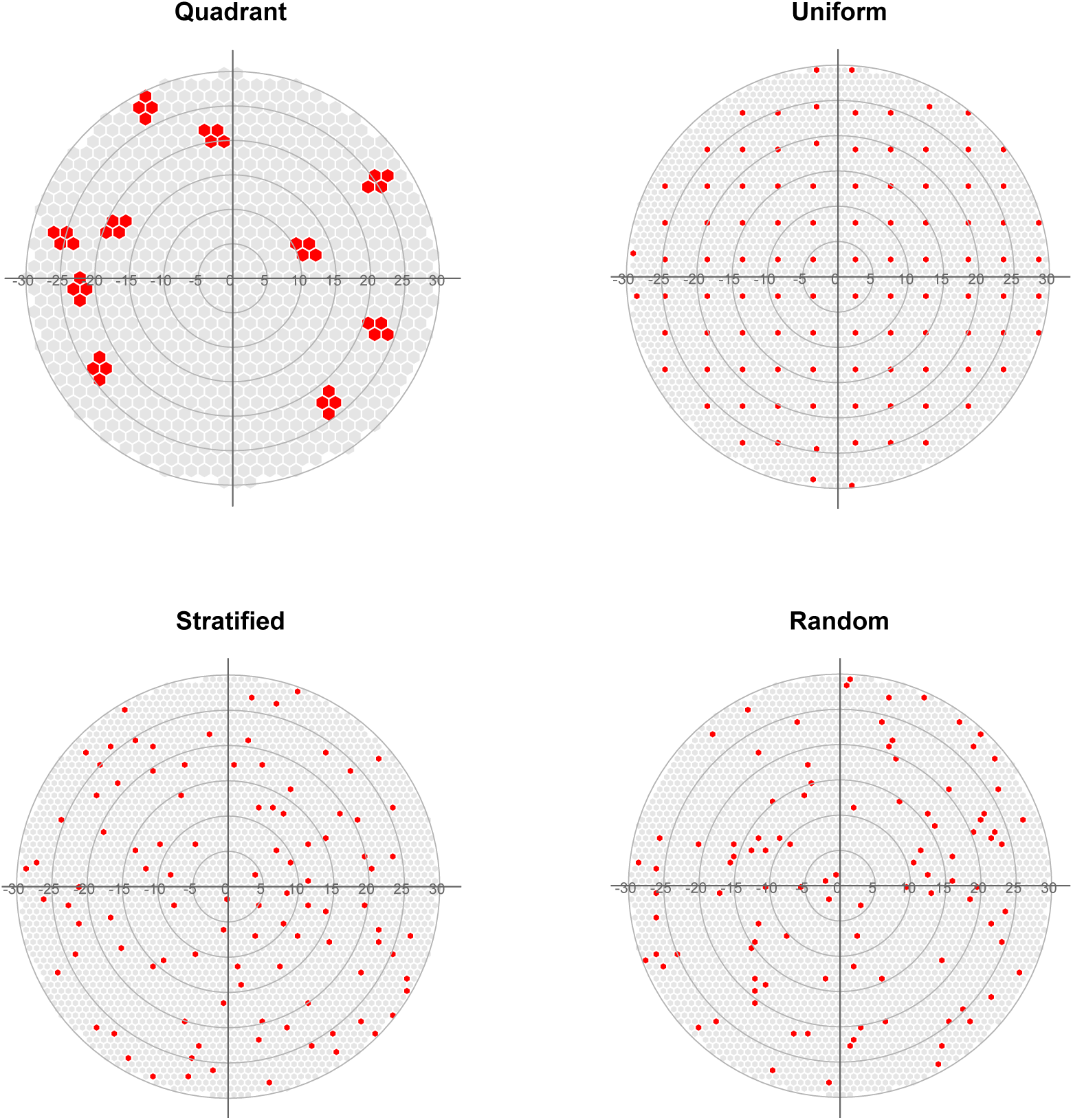
Examples of one simulation of each of the four under-roost sheet sampling designs explored in this study generated for a roost with a 30m radius. The quadrant design (top left), which follows methods found in previously published studies (Edson et al. 2015a, Field et al. 2011, 2015), is comprised of 10 3.6 × 2.6m sheets divided into 1.8 × 1.6m quadrants to produce 40 (10 × 4) quadrant-sized sheet areas for pooling urine samples. The other three designs (uniform, stratified, and random) are all ‘small-sheet’ designs that reduce sheet area, increase sheet number, and disperse sheets about the roost area. The small-sheet designs plotted above each contain 100 1m^2^ sheets. The stratified design is generated using a sequential inhibition process with and inhibitory radius of 2m.

### Calculating estimated prevalence

Given a roost area *A*, the polygons produced by the `sheetsamp` function (described above) generate the sheet sampling area *S*, so that *S* ⊂ *A*, and *S*_*h*_ = {*S*_1_, *S*_2_, *…, S*_*H*_}, where *H* is the total number of sampling sheets. We derived bat density from a simulated Poisson cluster point process and then estimated its intensity function *λ*(*x*) for area *A*. This method uses kernel density as an unbiased estimator of *λ*(*x*), which includes clustering of bats around trees, individual-level movement within the tree canopy, and roost-level movement to render 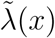. The expected number of bats roosting and moving above a specific sheet *S*_*h*_ placed at position (*x*_*h*_, *y*_*h*_) is the integral of the estimated intensity function 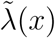 over the sheet area multiplied by the number of bats *N*_*b*_ generated by the stochastic point process.

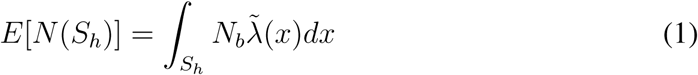

Bats in the upper strata of the canopy are less likely to contribute urine to the sheet below because of obstruction by individuals below or factors in the environment (e.g wind, tree branches). Therefore, a urine sample is collected from each of the sheets *S* according to a probability of urine contribution and collection *p*_*u*_, with variation given by *N* (*p*_*u*_, *σ*^2^). The number of individuals contributing to each pooled sample *C*_*b*_ is calculated as

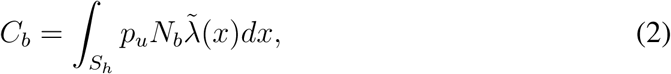

where *C*_*b*_ is a vector of length *H*, containing the number of contributing bats per sheet.

Assuming heterogeneous prevalence within the roost, the number of infected bats *D*_*b*_ in the sample is the sum of *C*_*b*_ independent Bernoulli trials with success probability equal to the true prevalence *p*.

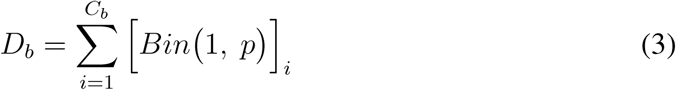

Given the number of infected bats *D*_*b*_ and the probability of urine collection *p*_*u*_, we can calculate the probability of obtaining a negative sheet as 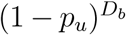. Assuming that urine contribution from one infected bat is sufficient to make a sheet sample positive, the infection status of all sheets is a binary vector *I*_*h*_ indicating the positivity for the *H* sheets of *S*.

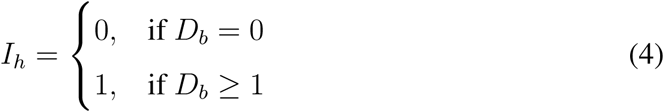

To calculate estimated sheet-level prevalence 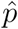, the number of positive sheets 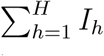 is divided by the number of urine samples collected at the roost *n*_*s*_, which is the sum of a binary vector indicating that the urine of more than one individual was contributed and collected for all of the *H* sheets of *S*.

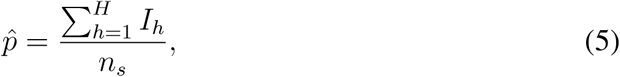

Where

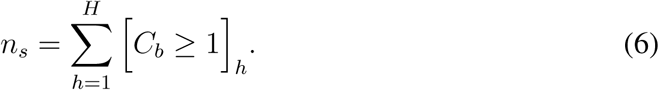

### Simulation scenarios

Each simulated iteration generates an estimated intensity function for bat density and then performs under-roost sampling using each of the four sampling designs. Therefore, each sampling design is tested using the same set of bat density functions, facilitating comparison. Parameters for sheet size *s* and number of sheets *H* were fixed for the quadrant-based design to replicate the previously implemented field methods described above. Parameters controlling sampling dimensions for the three small-sheet designs were either fixed or varied over a range of interest depending on the question the scenario was meant to address. A list of parameter values used in each scenario can be found in Table 1. For each iteration we calculated estimated prevalence (described above), along with additional analytic metrics such as the probability of obtaining a negative sheet 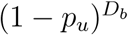, the occurrence of a false negative 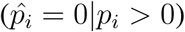, Moran’s I among sheets (Getis 1995), and the Clark-Evans R clustering coefficient for individual bat roosting positions (Clark and Evans 1954).

**Table 1:**
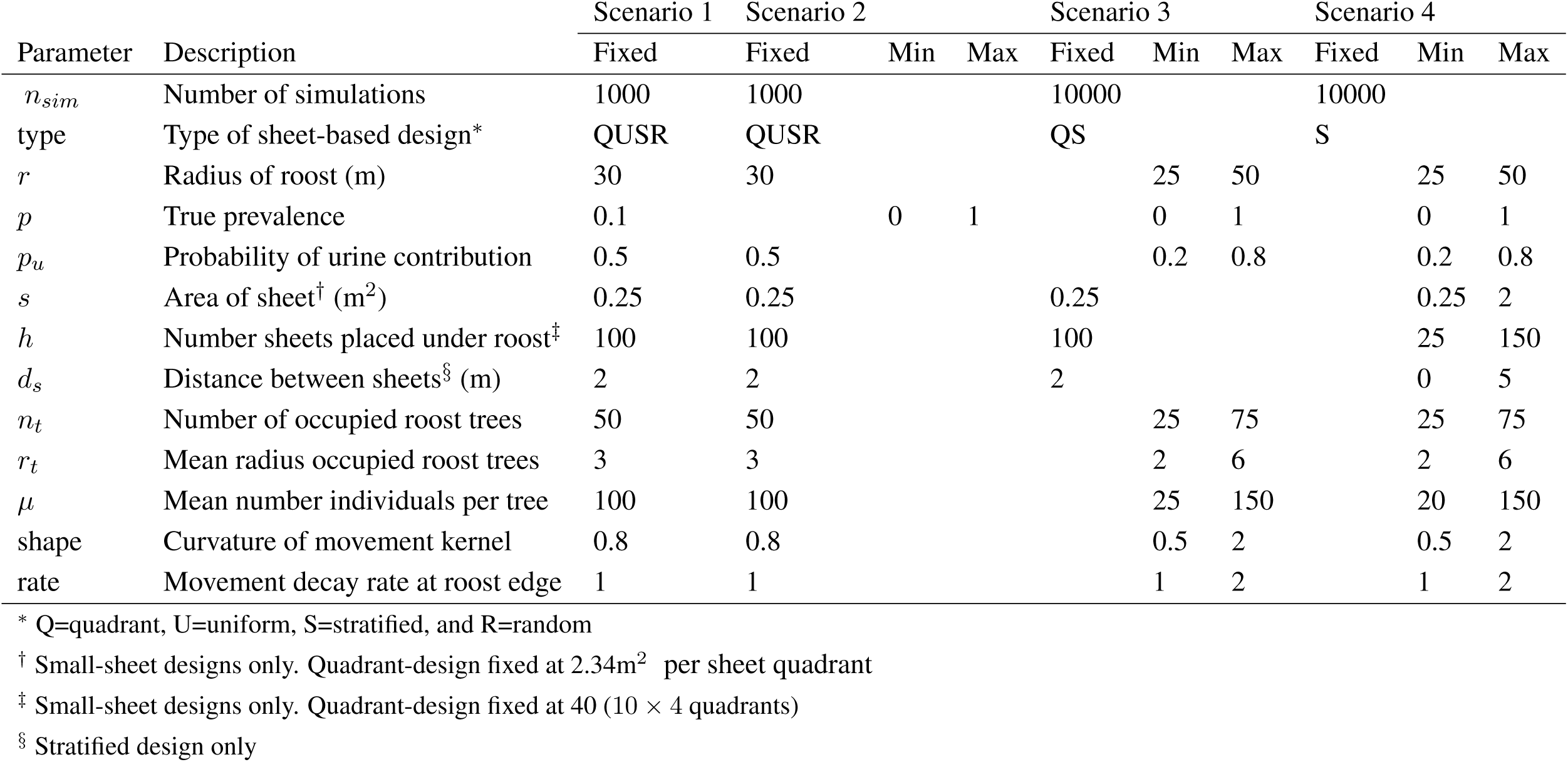
Fixed and varied parameter values used in each of the four scenarios. For scenarios 2-4, min and max set the minimum and maximum values of a uniform probability distribution within a random latin hypercube sampling approach.

In the first two scenarios we explored local sensitivity between estimated prevalence and some possible confounders and sources of bias, with values of most parameters fixed. To perform a simple comparison between the four under-roost sheet sampling methods, we fixed all values of bat density and movement to simulate a roost with a 30m radius and a mean number of 5000 individuals (see scenario 1 in Table 1). We performed 1000 simulations with true prevalence *p* set at a plausible value of 0.1. Estimated prevalence values were plotted, along with the probability of obtaining a negative sheet for each sampling design. To explore estimation bias over all values of true prevalence, we kept parameter values the same as simulation 1, but we allowed true prevalence to vary from 0 to 1, and then plotted true versus estimated prevalence along with mean estimation bias (scenario 2 in Table 1).

In scenarios 3 and 4, we used global sensitivity analysis, as described in Prowse et al. (2016), to identify the main sources of estimation bias and determine the optimal application of under-roost sheet sampling. Here, we performed a large number of simulations (*n*_*sims*_ = 10000), and allowed parameter values for each simulation to vary using latin hypercube sampling. We then analyzed the output using boosted regression trees (BRTs; De’ath 2007, Elith et al. 2008) as an emulator to link simulation inputs (varied parameters) with simulation outputs (we used estimation bias and false negative rate as responses). Parameter values were determined using the `randomLHS` function in the `lhs` package (Carnell 2016), and BRTs were fitted using the `gbm.step` function and the `gbm` and `dismo` packages (Hijmans et al. 2016, Ridgeway 2016). BRTs were fitted with appropriate error structure (Gaussian or Binomial) and meta-parameters set to ensure that the number of fitted trees exceeded 1000, following Elith et al. (2008), with tree complexity, learning rate, bagging fraction, and number of cross validation folds set to: 4, 0.005, 0.7, and 10 respectively. BRTs act as an effective emulator here because they fit complex non-linear relationships with up to third order interactions (tree complexity=4) among model parameters. Relative variable influence and individual response curves for each variable further allow general description of how sensitive estimation bias is to each parameter.

In scenario 3, we compare the quadrant-based design with the stratified design while accounting for the variability in all other parameters to determine the main drivers causing differences in estimation bias. We chose to use only the stratified design as a candidate small-sheet design because the first two simulations suggested that the three small-sheet designs produce similar results, and the stratified design is most plausibly replicated in the field. Based on preliminary models, it appeared that a small-sheet sampling design which used ∼100 sheets with an area of ≤ 1 × 1m^2^ could attain low estimation bias. So, we fixed the parameters controlling sheet dimensions accordingly to facilitate comparison between the quadrant and stratified methods (see simulation 3 in Table 1).

To explore the optimal application of the stratified sampling design, we performed a global sensitivity analysis using only the stratified sampling design in scenario 4. All parameters were varied as in scenario 3, however sheet area *s*, number of sheets *H*, and distance between sheets (*d*_*s*_; previously fixed at 2m) were also varied over intervals of interest (scenario 4 in Table 1). We used a latin hypercube to sample the parameter space, and then fitted two BRT models using the variables that control the sheet sampling design as predictors (i.e. sheet area, number of sheets, distance between sheets, and number of samples). The first model we fitted with Gaussian error and estimation bias as the response, and the second with Binomial error and a binary response indicating occurrence of a false negative prediction for viral presence.

## Results

When we compared the quadrant-based sheet design to the small-sheet designs with fixed model parameters (scenario 1 in Table 1), we found that at a low value of true prevalence (0.1) the quadrant design exhibited strong positive bias and all three small-sheet designs produced similar estimates close to the fixed value of true prevalence (see top row of Figure 3). The differences in estimated values can be partially attributed to the increased number of bats that roost and move above the larger sheets, which decrease the probability of obtaining a negative sheet (see bottom row of Figure 3). Local sensitivity analysis revealed that, at a low value of true prevalence, prevalence estimation for the quadrant-based design is sensitive to spatial auto-correlation among sheets (Moran’s I) and clustering of bat roosting positions (Clark-Evans R) (Figures S2 and S3). However, the small-sheet designs are sensitive to the number of bats in the roost (*N*_*b*_) (Figure S4). This indicates that, at low values of individual-level prevalence, the quadrant based method remains sensitive to viral presence regardless of the roost population size, but will tend to over-estimate viral prevalence due to the spatial clustering of individuals common to most tree roosting bats. Conversely, small-sheet methods appear less affected by clustering and spatial auto-correlation among sheets, but they are likely to be less sensitive to viral presence at low population sizes.

**Figure 3:**
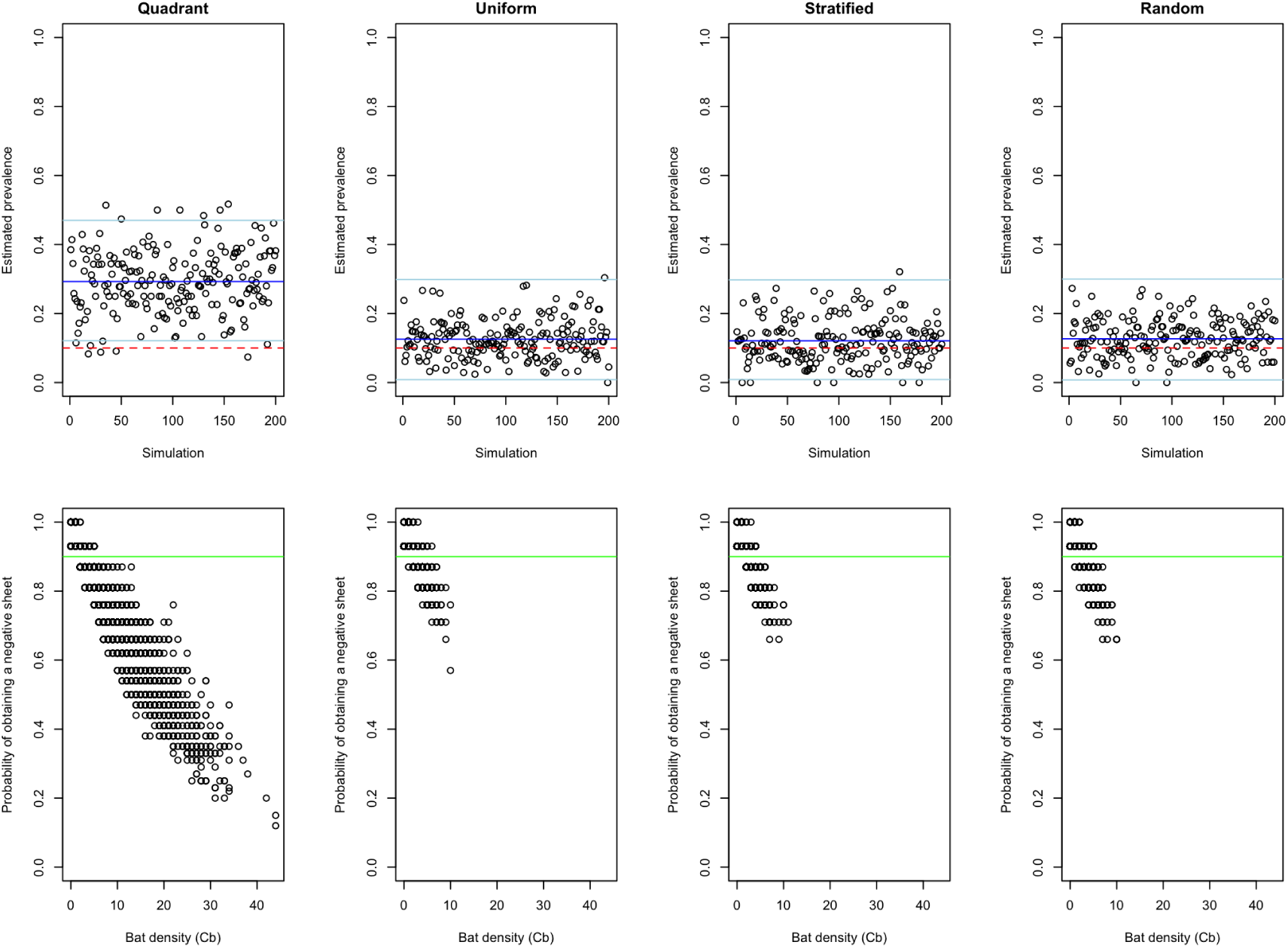
Results of scenario 1, where all parameters are fixed to facilitate comparison among the four sampling designs (Table 1). Top row displays the estimated prevalence of each simulation, with the mean of all simulations indicated by the blue line and the mean error due to sample size indicated by the light blue lines. The dashed red line indicates the fixed value of true prevalence (0.1). Bottom row displays the probability of obtaining a negative sheet (1 – *p*_*u*_)^*D*^_*b*_ as a function of the number of bats contributing urine to each sheet *C*_*b*_. The green line indicates the expected probability of obtaining a negative sheet given by (1 – *p*).

In scenario 2, where we allowed true prevalence to vary between 0 and 1 (Table 1), we found that the quadrant design had 5–7 times the positive bias as the small-sheet designs. The mean estimation bias was 0.21 for the quadrant design, and 0.4, 0.3, and 0.4 for the uniform, stratified, and random designs respectively. This suggests that, for a roost size of 3000–8000, the estimation bias will consistently be greater for the quadrant design, especially for intermediate values of individual-level prevalence. Additionally, the similarity among the uniform, stratified, and random designs indicates that the exact spatial pattern of the small-sheet method is not important—estimation bias is improved by reducing sheet size, increasing the number of sheets, and spreading sheets out within the roost area.

Scenario 3 showed significant differences in estimation bias between quadrant and stratified designs, even when we allowed all parameters to vary (Figure 5e). Summary of simulation output with the BRT emulator showed higher bias for the quadrant design, which is most strongly influenced by the total number of individual bats sampled across all sheets(∑ *C*_*b*_; Figures 5a and b). This suggests that the larger sheet area in the quadrant design allows pooling of urine samples from more individuals, making the prevalence estimates more sensitive to increases in population size. Further, a quadrant-based design allows up to four ‘independent’ pooled samples to be adjacent each other, effectively inflating the number of positive sheets, illustrated by higher estimated prevalence associated with high values of Moran’s I in Figure 5d. In general, both sampling designs are positively influenced by intermediate values of true prevalence, number of bats in the roost (leading to a greater number of total bats contributing to each sample), and spatial auto-correlation among sheets. However, the influence of these factors is diminished in the stratified design, as shown by the orange points in Figures 5b–f.

**Figure 4:**
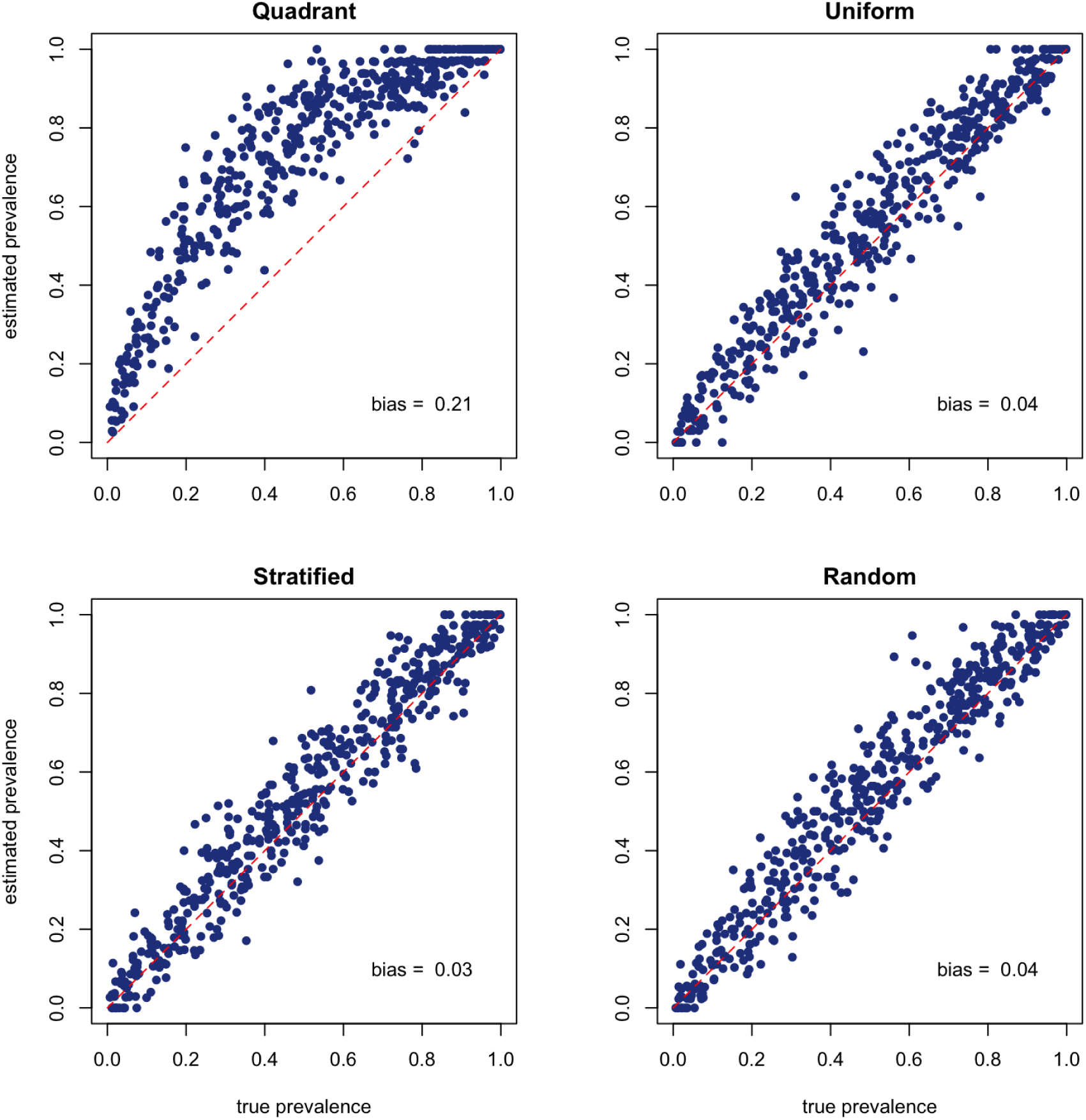
Results of 1000 simulations performed over all possible values of true prevalence for four different under-roost sheet sampling designs (see scenario 2 in Table 1). The dashed red line indicates 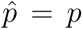, and mean estimation bias for all simulations is printed in the lower right corner of each plot.

**Figure 5:**
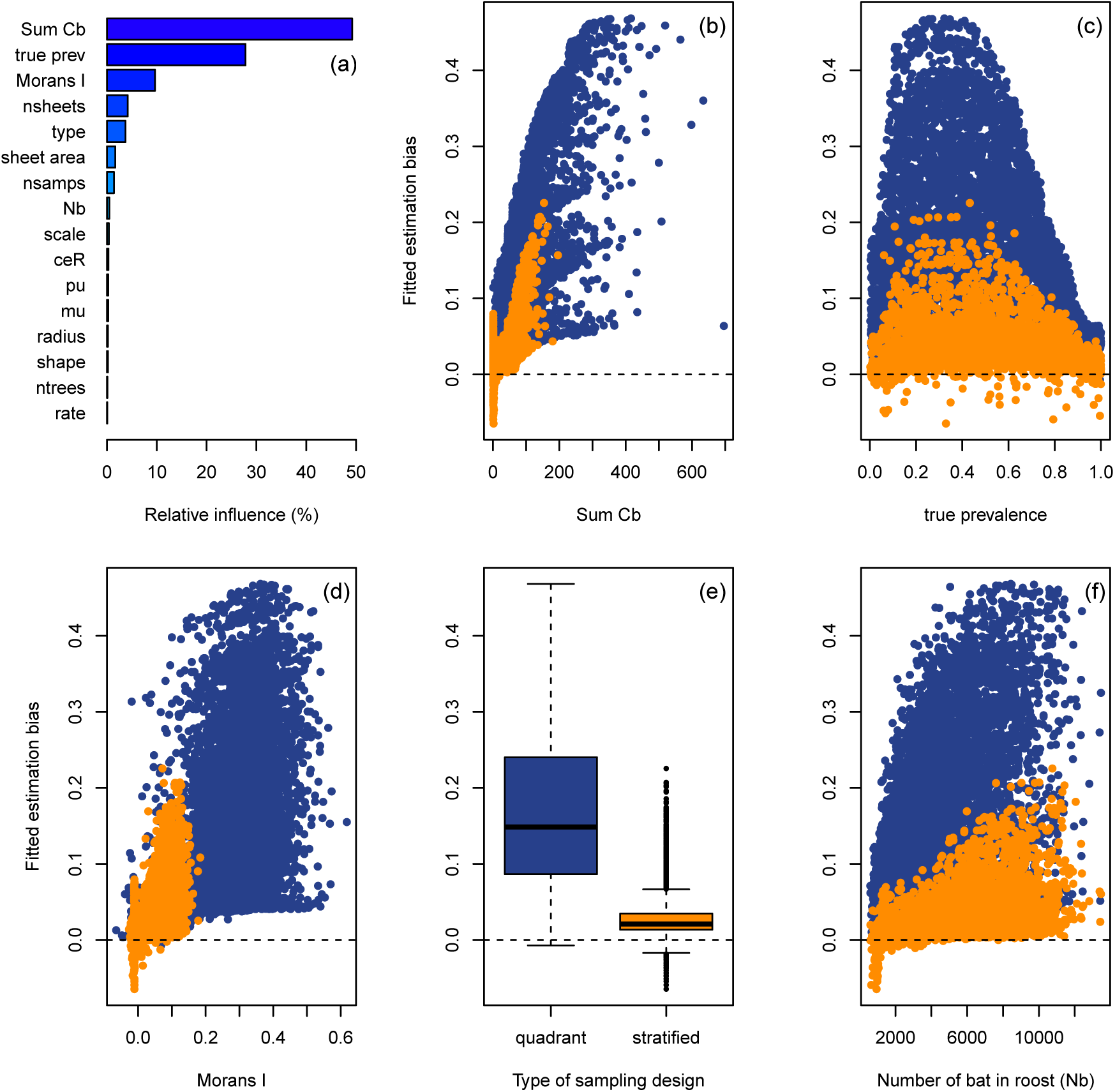
Results of the global sensitivity analysis performed in scenario 3, where the quadrant (blue points) and stratified (orange points) designs are compared to determine what drives differences in estimation bias between the two designs. Table 1 shows the parameters used in the simulation. The barplot (a) shows the relative influence of each parameter determined by a boosted regression tree emulator. Plots e–f show the value of estimation bias fitted by the emulator as a function of five influential parameters.

When we further explored the influence of sheet dimensions for the stratified design (scenario 4 in Table 1), we found that sheet area *s* and number of samples collected *n*_*s*_ influenced estimation bias and probability of false negatives the most, and the number of sheets *H* and distance between sheets *d*_*s*_ had less influence (Figure 6). Specifically, estimation bias increases for sheet area greater than 0.5m^2^, but the probability of false negatives increases for sheet area less than 0.75m^2^. Suggesting that sheet areas in the range of 0.5–1m^2^ would provide a balance of the two sources of sampling bias (Figures 6a and e). The number of sheets had no influence on estimation bias, however, sampling designs with less than 80 sheets had higher probability of false negatives (Figures 6b and f). Minimum distance between sheets did not have a significant effect on either source of sampling bias, however, distances between 2–3m fitted the lowest maximum probability of false negatives (Figures 6b and f). The number of samples collected *n*_*s*_ exhibited the largest influence among sheet dimension parameters. Estimation bias increased with a larger number of collected samples, with the possibility for underestimation when under 20 samples were obtained (Figure 6d), and the probability of false negatives increased below 30–40 samples (Figure 6h). In general, these results indicate that collecting 30–40 pooled urine samples with a stratified sheet sampling design that uses 80–100 sheets, each with an area of 0.5–1m^2^, that are separated by a minimum distance of 2–3m, would provide optimal application of the under-roost sampling technique that minimizes error introduced by estimation bias and false negatives. Further, we calculated the proportion of simulations matching the parameters stated above and found that, given a roost population size greater than 5000, 89% of simulations had at least 30 sheets that collected a urine sample, and 64% collected at least 40 samples (Figure S5).

**Figure 6:**
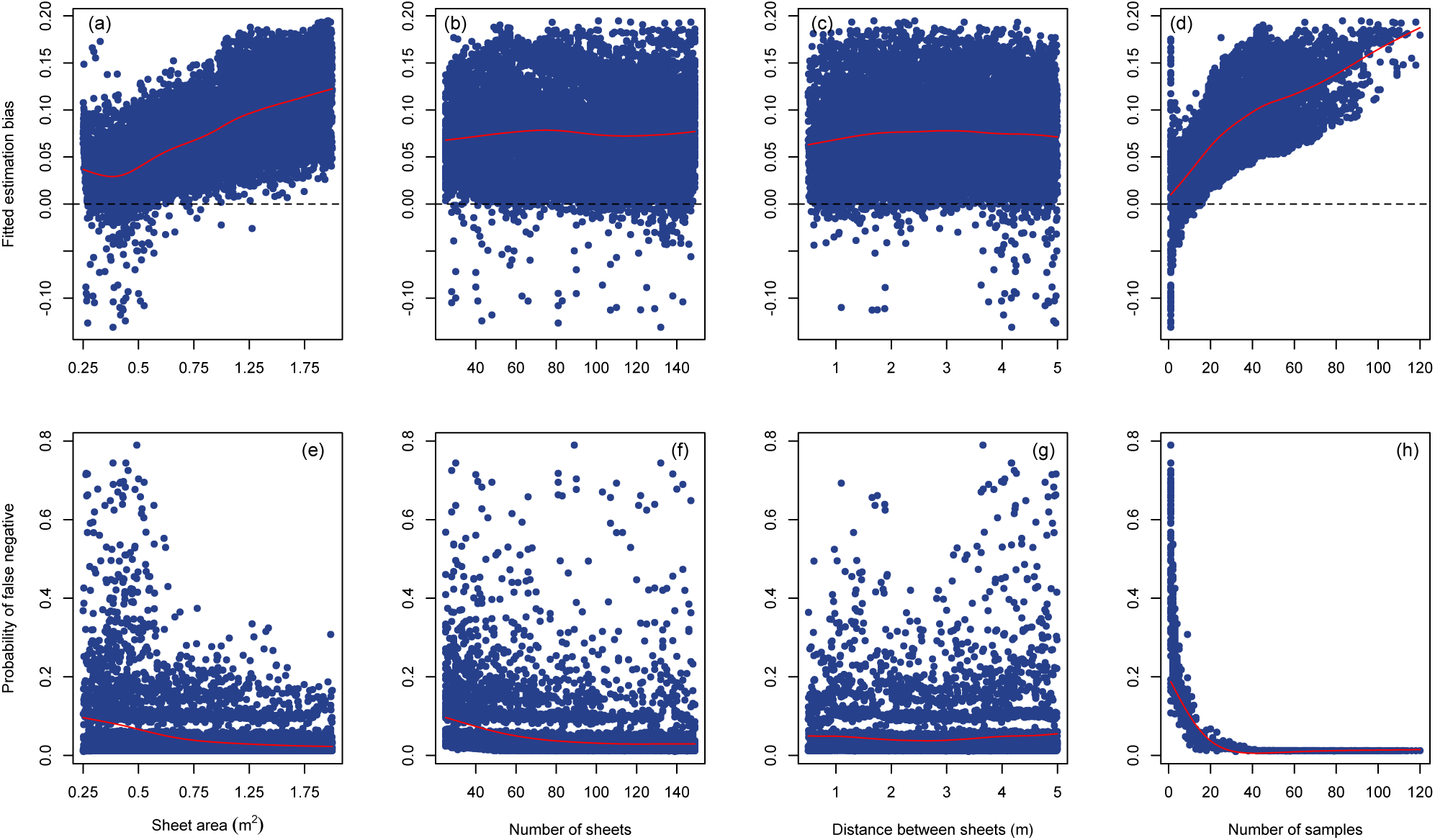
Global sensitivity analysis of scenario 4, where the influence of sheet dimension parameters are explored to determine optimal application of the stratified sheet sampling design. The plots display results from two boosted regression tree emulators: one for estimation bias (top row), and the other for the probability of false negatives (bottom row). Each response is plotted against sheet dimension parameters (from left to right): sheet area *s*, number of sheets *h*, minimum distance between sheets *d*_*s*_, and number of samples collected *n*_*s*_. The red lines indicate the trend of the points given by smooth spline regression (`sreg` function in the `fields` R package; Nychka et al. (2015))

## Discussion

Under-roost sampling of bat viruses has been employed previously in Africa, Asia, and Australia, however little attention has been given to the effects of sampling bias or optimization of sampling designs. We present the first modeling study to theoretically investigate under-roost sampling in detail. The simulation scenarios we developed enable inference on the relationship between individual-level prevalence and roost-level prevalence estimated for a generic population of tree roosting bats. Specifically, our results provide three key insights that will help to refine the application of under-roost sampling in the surveillance of infectious viruses in wild bat populations. First, sampling designs which use large sheets (larger than ∼1m^2^), and/or sheet-quadrants to pool urine samples are sensitive to viral presence, but they potentially over-estimate viral prevalence with a bias up to 7 times greater than a design with a greater number of smaller sampling sheets (Figure 4). Second, estimation bias is affected by the number of individuals allowed to contribute to a pooled sample and spatial auto-correlation among sampling sheets, however these sources of bias can be reduced by adjusting the sheet sampling design (Figure 5). And third, assuming a roost population size of over 5000, estimation bias can be sufficiently reduced by collecting 30–40 pooled urine samples using a stratified sheet sampling design that uses 80–100 sheets, each with an area of 0.75–1m^2^, that are separated by 1–3m (Figures 6 and S5). Our insights from simulation models provide well-informed hypotheses about the optimal sheet design for under-roost sampling, which facilitates further development within a model-guided fieldwork approach (Restif et al. 2012).

Our recommendations to optimize under-roost sampling differ from those previously implemented in the field in that they reduce the size of sheet area, increase the number of sheets, and disperse them about the roost area. In relation to the best-described methods in the literature, this is roughly equivalent to halving the size of sheet quadrants in Edson et al. (2015a) and Field et al. (2015) to make 80 0.9 × 0.8m sheets, and then separating each of them by 1–3m. Or relative to Wacharapluesadee et al. (2010), the sheets could remain 1.5 × 1.5m (or be reduced to 1 × 1m), but the total number of sheets could be increased by 3–4 times. However, we acknowledge that our recommendations are derived from simulation models that generalize a broad array of roost areas and population sizes that do not take into account local topography around a roost. Local factors at the roosting site (e.g. physical obstructions, understory vegetation, slope) must be considered when applying sampling designs in the field. Further, ‘optimal’ application of an under-roost sampling design is still inherently limited to pooled sheet-level estimates of prevalence. As our results show, this makes it difficult to entirely remove positive bias associated with such data aggregation, however it can be mitigated with a sheet design that reduces the area of urine pooling and limits spatial auto-correlation among sheets.

We hypothesize that under-roost sampling designs as they have been applied in the past are poorly suited to studying viral dynamics because of positive sampling bias. For example, Páez et al. (2017) analyzed data from an under-roost sampling study (Field et al. 2015), and noted that a large amount of variation in viral prevalence was explained by differences in sampling sheets, indicating that population structure within roosts or sampling bias may have introduced additional variation in estimated prevalence. In light of the results from our simulation models, pooling urine samples drawn from large sheet areas effectively inflates the number of Bernoulli trials in each Binomial sample. This may be observed as overestimation when the pooled samples are subsequently used to calculate prevalence in such studies. Therefore, collecting pooled urine samples from a smaller sheet area may reduce the number of bats contributing to a sample and the potential for overestimation, with the caveat that smaller sheets are less likely to collect urine samples, necessitating a larger number of sheets placed under the roost.

We have shown that sheet design in under-roost sampling can have a significant impact on both the estimation of viral prevalence and the false negative rate when determining viral presence. The sampling design employed, therefore, depends on the aim of the study, because viral discovery and studies on dynamics require different approaches. Research focusing on viral discovery requires field methods that reduce the probability of a false negative in regard to viral presence (sensitivity). Studies on dynamics must estimate prevalence with low bias, requiring samples that are accurately classified as present and absent (specificity). Further, the volume of urine sample required by the diagnostic test will determine how large the sheet area must be when pooling urine samples. For instance, if you are only interested in the presence or absence of viral RNA in a sample, RT-PCR requires a mere 50-150*µ*L sample, allowing a few droplets from a rather contained area to be taken. If however, a larger volume is required for sequencing or multiple assays, then up to 1–2mL may be required, necessitating a larger pooled sample from a greater area that is more susceptible to bias associated with data aggregation (Robinson 2009). Therefore, if a study includes multiple aims, an efficient adaptation of a small-sheet design includes pooling urine over multiple spatial scales, with samples pooled over a large area to test for viral presence with high sensitivity *and* samples pooled over a small area for estimating individual-level prevalence with high specificity. For example, a researcher might put down 100 1 × 1m sheets, and collect 40 100 *µ*L small pooled samples from 30–40 separate sheets. The remaining urine can be pooled across multiple sheets to form larger pooled samples that provide higher sensitivity to viral presence. This approach is similar to the aforementioned herd-level testing in veterinary epidemiology (Christensen and Gardner 2000), where a herd of livestock is first tested by pooling multiple samples as a low-cost test with high sensitivity. If virus is found in the large-scale pooled samples, then many the small-scale pooled samples can be used to accurately estimate prevalence.

Our simulation models and recommendations for a small-sheet sampling design provide an important contribution that facilitates future research. Specifically, we propose that under-roost sampling can be further developed with two important avenues of research: i) a comparative field study to quantify differences in sheet sampling designs in a model-guided field work approach (Restif et al. 2012), and ii) modeling studies that incorporate previous work on estimating individual-level prevalence from pooled samples (Cowling et al. 1999, Hauck 1991) to investigate bias correction for existing and future field data. Given the challenges associated with under-roost sampling, it remains an attractive supplement to catching and sampling individual bats. If applied in a manner suited for study aims, it can achieve longitudinal sampling of a bat population at the roost-scale that is both cost effective and reduces exposure to infectious viruses. Further development of the sampling technique into a replicable sampling method is also advantageous, because it enables population level surveillance of infectious viruses in bats, which provide insights into ecological processes that drive spillover and emergence of bat-borne viruses over large spatial scales.

## Acknowledgments

JRG conducted the research with support from an Australian Government Research Training Program Scholarship.

## Supplementary Material

**Figure S1:**
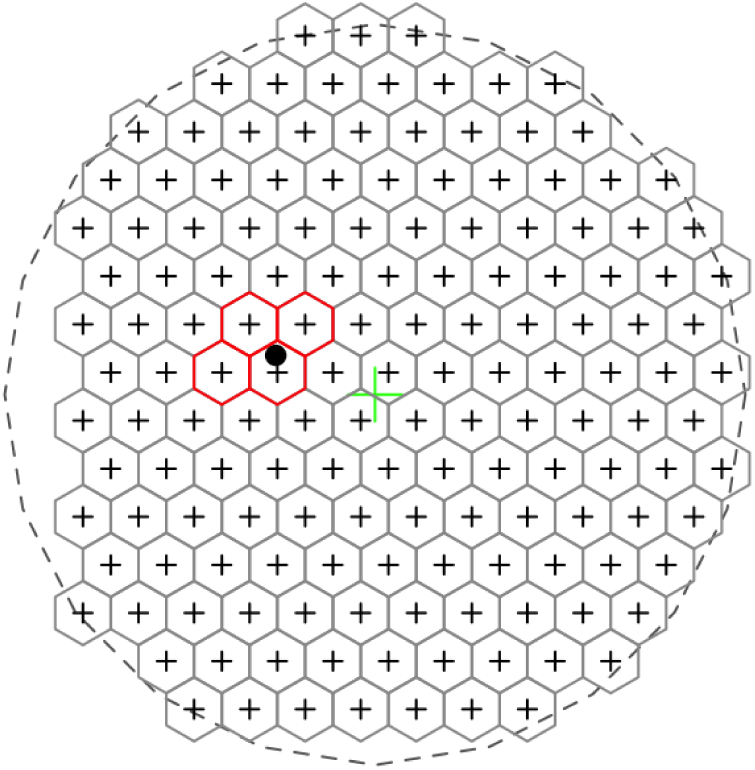
Example of construction how sheet areas are defined using the quadrant-based under-roost sheet sampling technique. The schematic shows a grid of hexagonal tiles filling a circular roost area. Cell centroids are marked with a black cross. One large sheet with four quadrants is made by selecting a sheet location (black point) and then selecting the four nearest centroids.

**Figure S2:**
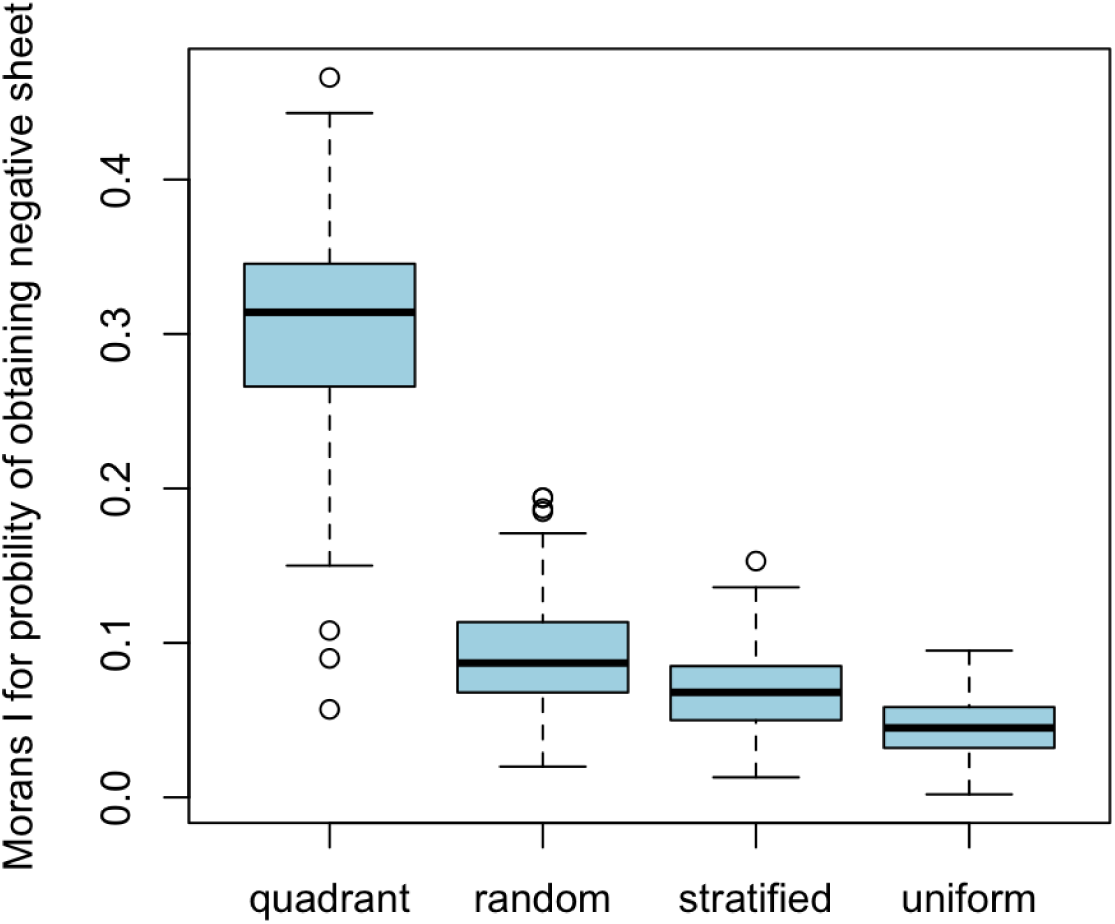
Boxplots showing the variation in Moran’s I calculated as part of the local sensitivity analysis in Simulation 1. The amount of spatial autocorrelation in the probability of obtaining a negative sheet is shown on the y-axis, and the four under-roost sheet sampling designs on the x-axis.

**Figure S3:**
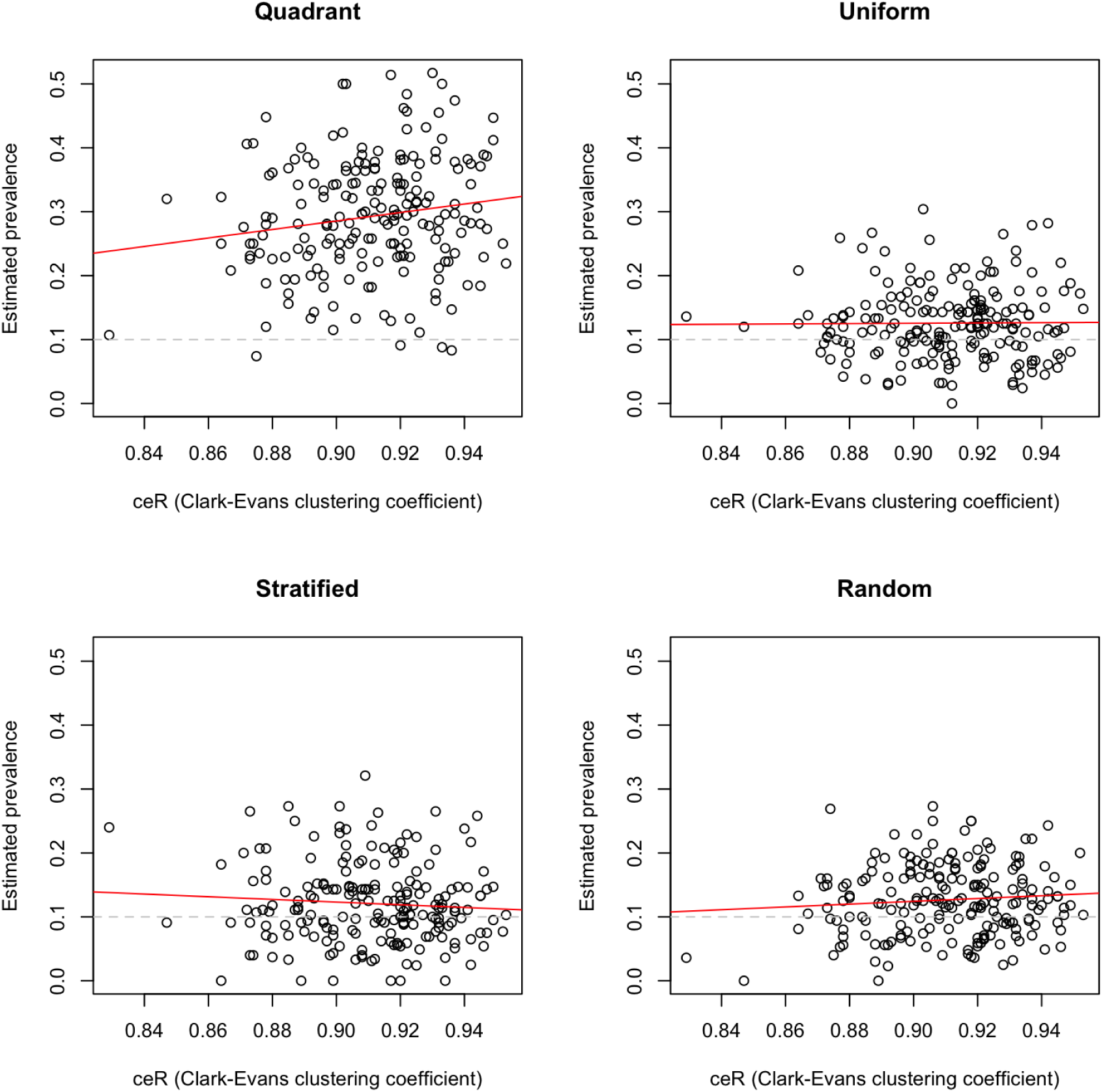
Scatterplots showing the variation in the Clark-Evans R clustering coefficient calculated as part of the local sensitivity analysis in scenario 1. The Clark-Evans R gives a measure of how clustered bat roosting positions are within the simulated roost. For each of the four sheet sampling designs, the estimated values of viral prevalence 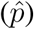 is plotted on the y-axis, and the Clark-Evans R (ceR) is plotted on the x-axis. Linear model trend lines are shown in red and the value of true prevalence (*p*) set in the local sensitivity analysis is the dashed gray line.

**Figure S4:**
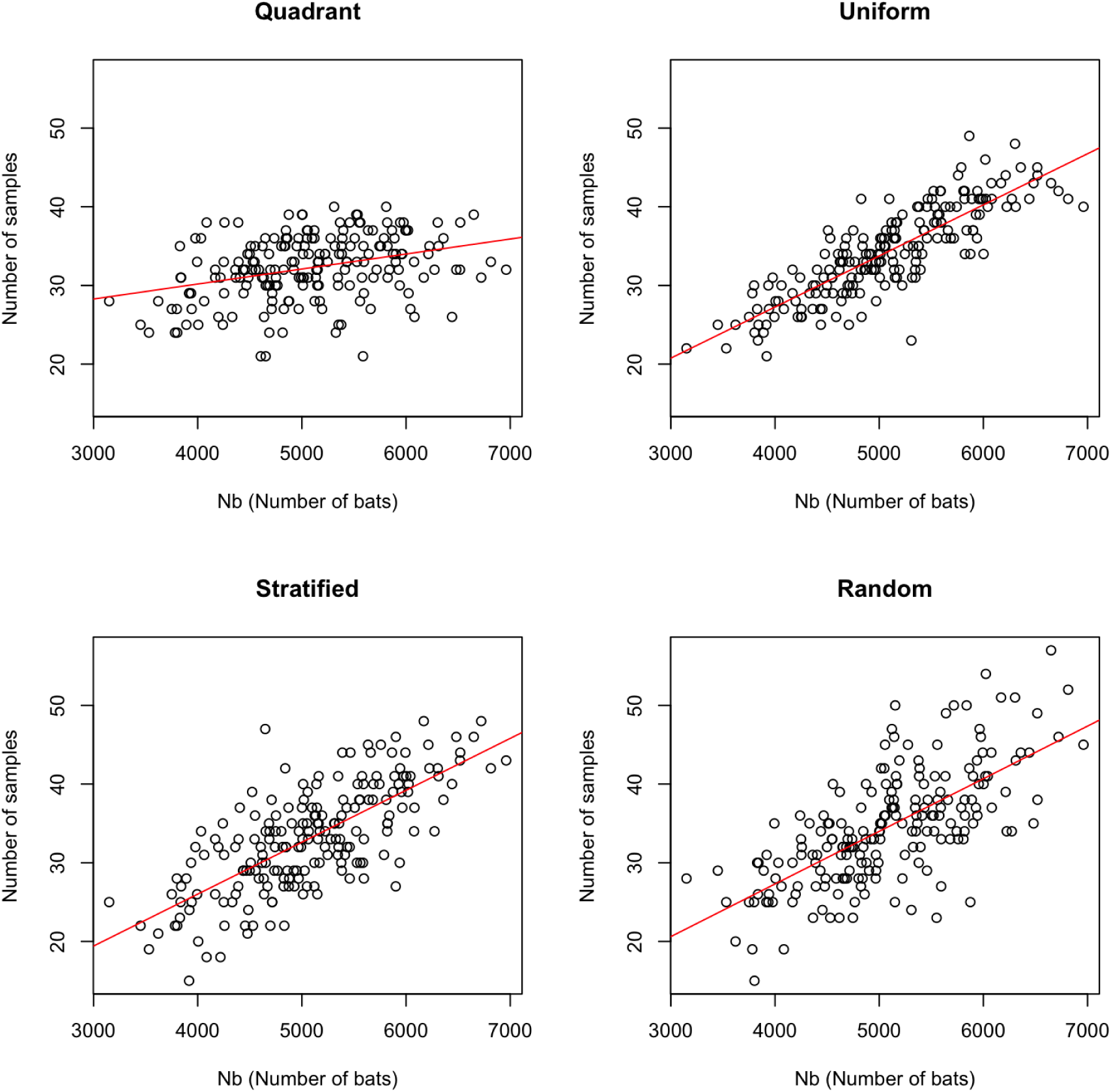
Scatterplots showing the variation in the number of total bats in the roost (*N*_*b*_) calculated as part of the local sensitivity analysis in scenario 1. For each of the four sheet sampling designs, the estimated values of viral prevalence 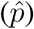 is plotted on the y-axis, and the number of bats (*N*_*b*_) is plotted on the x-axis. Linear model trend lines are shown in red and the value of true prevalence (*p*) set in the local sensitivity analysis is the dashed gray line.

**Figure S5:**
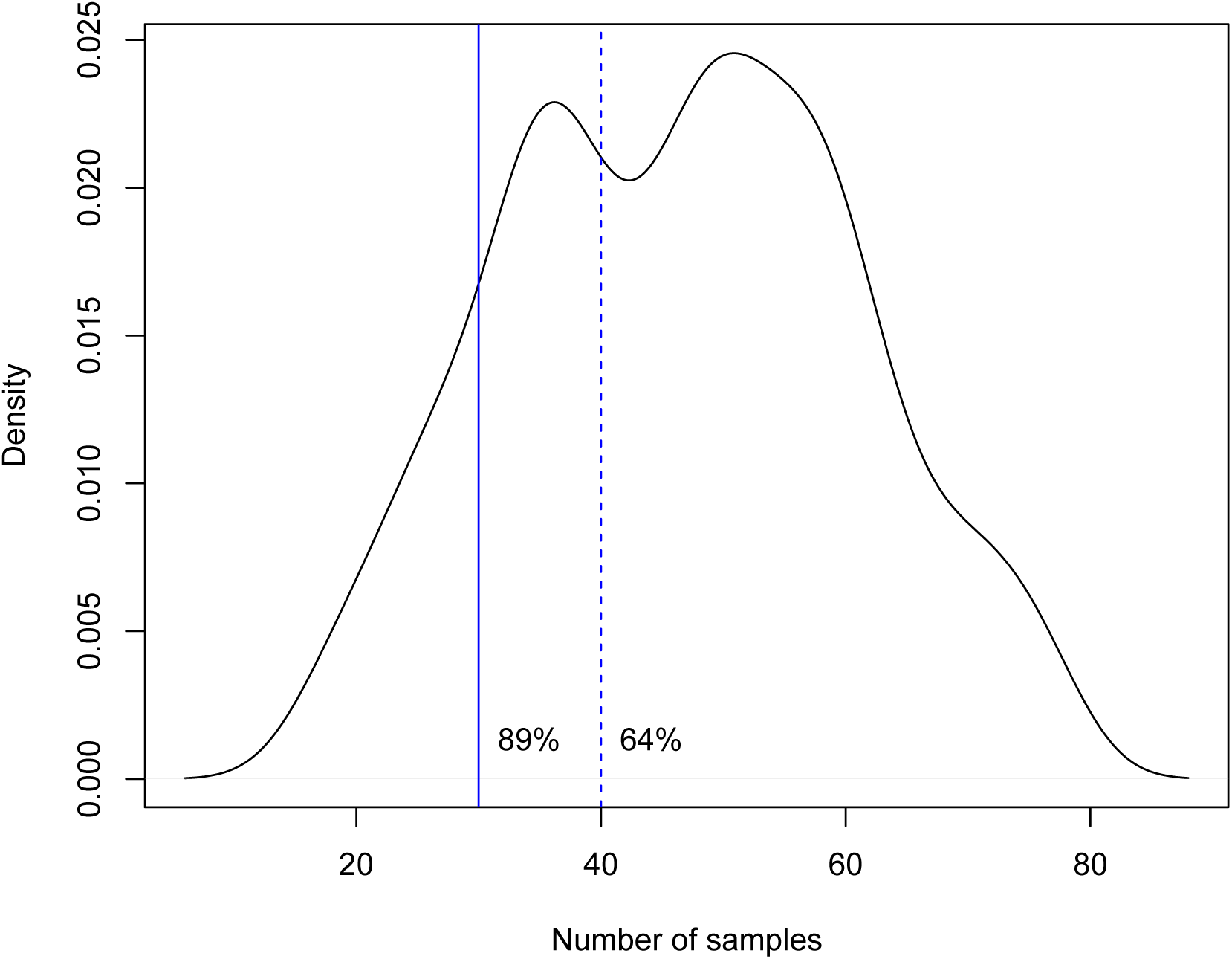
Distribution of the number of samples collected for simulations that use a stratified sheet sampling design at a roost of *>* 5000 individuals, where the number of sheets *n*_*s*_ is 80–100, the area of the sheets *s* is 0.75–1m^2^, and the distance between the sheets is 1–3m. Based on our results, 89% of simulations had at least 30 sheets that collected a urine sample, and 64% that collected at least 40 samples.

